# Single neurons throughout human memory regions phase-lock to hippocampal theta

**DOI:** 10.1101/2020.06.30.180174

**Authors:** Daniel R. Schonhaut, Ashwin G. Ramayya, Ethan A. Solomon, Nora A. Herweg, Itzhak Fried, Michael J. Kahana

**Affiliations:** Department of Neuroscience, Perelman School of Medicine, University of Pennsylvania, Philadelphia, PA 19104, USA; Department of Neurosurgery, Hospital of the University of Pennsylvania, Philadelphia, PA 19104, USA; Department of Bioengineering, University of Pennsylvania, Philadelphia, PA 19104, USA; Department of Psychology, University of Pennsylvania, Philadelphia, PA 19104, USA; Department of Neurosurgery, David Geffen School of Medicine and Semel Institute for Neuroscience and Human Behavior, University of California, Los Angeles, CA 90095, USA; Tel-Aviv Medical Center and Sackler Faculty of Medicine, Tel-Aviv University, Tel-Aviv 69978, Israel

## Abstract

Functional interactions between the hippocampus and cortex are critical for episodic memory. Neural oscillations are believed to coordinate these interactions, and in rodents, prefrontal neurons phase-lock to hippocampal theta oscillations during memory-guided behavior. We assessed inter-regional phase-locking to hippocampal oscillations in humans by recording 1,233 cortical and amygdala neurons and simultaneous hippocampal local field potentials in 18 neurosurgical patients. We identified 362 neurons (29.4%) from multiple regions that phase-locked to rhythmic hippocampal activity, predominantly at theta (2-8Hz) frequencies. Compared to baseline spiking, strong theta phase-locking coincided with regionally-specific increases in hippocampal theta power, local and hippocampal high frequency activity, and cross-frequency power correlations between the hippocampus and a phase-locked neuron’s local region. These results reveal that spike-time synchrony with hippocampal theta is a defining feature of cortico-hippocampal functional connections in humans. We propose that theta phase-locking could mediate flexible inter-regional communication to shape the content and quality of episodic memories.

## Introduction

The hippocampus is the operational hub of the episodic memory system, a spatially distributed brain network that enables us to remember past experiences in rich detail, together with the space and time in which they occurred (Eichenbaum, 2000; Moscovitch et al., 2016). To serve in this capacity, the hippocampus must maintain precise but flexible functional connections with the rest of the memory system. Understanding the mechanisms that govern these connections is of fundamental interest to systems neuroscience, and progress in this area could accelerate efforts to develop treatments for memory disorders and age-related memory decline.

A leading hypothesis is that hippocampal theta (2-8Hz) oscillations facilitate its interactions with other brain regions (Buzsáki, 2010; Fell and Axmacher, 2011; Moscovitch et al., 2016). Hippocampal neurons are more receptive to synaptic excitation at specific theta phases (Kamondi et al., 1998), so well-timed inputs can more effectively drive activity than inputs at random phases (Fries, 2005). Long-term potentiation and long-term depression in the rodent hippocampus are also theta phase-dependent (Hyman et al., 2003), providing a putative link between the phase at which inputs arrive and how strongly they are encoded into memory. Experimental evidence for this hypothesis comes largely from studies in the rat medial prefrontal cortex (mPFC), a downstream target of hippocampal area CA1. mPFC neurons phase-lock to hippocampal theta during short-term memory tasks (Siapas et al., 2005; Hyman et al., 2005; Sirota et al., 2008), and stronger phase-locking predicts better performance (Jones and Wilson, 2005; Hy-man et al., 2010; Benchenane et al., 2010; Fujisawa and Buzsáki, 2011) and greater information transfer between mPFC and hippocampal neurons (Ito et al., 2018; Padilla-Coreano et al., 2019). Phase-locking to hippocampal theta is also prevalent among cells in many other regions, including the entorhinal cortex (EC), amygdala, parietal cortex, thalamic nucleus reuniens, and some subcortical and brainstem nuclei (Kocsis and Vertes, 1992; Sirota et al., 2008; Fujisawa and Buzsáki, 2011; Bienvenu et al., 2012; Fernández-Ruiz et al., 2017; Ito et al., 2018). Theta phase-synchronization could thus be a general mechanism for relaying information between the hippocampus and a broad network of memory-related regions.

In humans, macroelectrode local field potential (LFP) recordings in epilepsy patients have revealed sporadically-occurring theta oscillations in the hippocampus and cortex during virtual navigation and episodic memory engagement (Fell et al., 2011; Watrous et al., 2013; M. Aghajan et al., 2017; Bohbot et al., 2017). Studies relating theta power to memory have produced equivocal results (Herweg et al., 2020), but macroscale theta phase-synchronization within the medial temporal lobe (MTL) and PFC has consistently correlated with better memory performance (Babiloni et al., 2009; Watrous et al., 2013; Solomon et al., 2017; Zheng et al., 2019; Kunz et al., 2019). Considerably less is known about how oscillations relate to single-neuron activity in humans than in rodents. An early study in epilepsy patients found that a large percentage of MTL and neocortical neurons phase-locked to theta (among other frequency) oscillations in their local vicinity as subjects navigated through a virtual environment (Jacobs et al., 2007), and another study discovered MTL neurons phase-locked more strongly to locally-recorded theta oscillations while subjects viewed images that they later recognized than those that they forgot (Rutishauser et al., 2010). These findings indicate that neural activity within the human episodic memory system is organized in part by a theta phase code. However, single-neuron phase-locking to remotely-recorded oscillations has not been identified in humans, and the hypothesis that hippocampal theta oscillations facilitate communication with other brain regions therefore lacks an important element of empirical validation.

Here, we leveraged the rare opportunity to record single neurons simultaneously with hippocampal LFPs in subjects with pharmacologically intractable epilepsy. We identified a substantial fraction of neurons that fired phase-synchronously with hippocampal theta oscillations. Phase-locking rates were greatest in regions structurally connected to the hippocampus, but significant neocortical and contralateral phase-locking suggests that neurons can synchronize with hippocampal theta oscillations across polysynaptic distances. Lastly, we observed pronounced changes in LFP power when neurons fired near their preferred hippocampal theta phase (“strong phase-locking”), relative to baseline spiking. Strong phase-locking coincided with increased hippocampal theta power, greater local and hippocampal high frequency activity (HFA), and more correlated neural activity patterns (LFP power across frequencies) between the hippocampus and a phase-locked neuron’s local region. These effects were specific to the hippocampus and the phase-locking region, supporting an ability for hippocampal theta oscillations to coordinate regionally-precise patterns of cortico-hippocampal connectivity in the human brain.

## Results

We recorded hippocampal micro-LFPs and extracellular spikes from 1,233 neurons in the EC, amygdala, parahippocampal gyrus, orbitofrontal cortex, anterior cingulate, and more sparsely sampled neocortical regions as subjects (18 neurosurgical patients, 1-4 sessions per subject) played a virtual navigation game on a laptop computer (Table 1) (Ekstrom et al., 2003). We did not compare physiology to behavioral measures in this study (see Discussion). Neural firing rates were log-normally distributed (median = 2.3Hz), and we recorded between 11 and 149 (mean=68.5) neurons per subject.

**Table 1:** Neuron locations. Rows show how many subjects, testing sessions, and neurons were recorded from each brain region.

| Neuron region | Subjects | Sessions | Neurons |
| --- | --- | --- | --- |
| Amygdala | 14 | 30 | 327 |
| Anterior cingulate | 5 | 12 | 101 |
| Entorhinal cortex | 12 | 30 | 221 |
| Orbitofrontal cortex | 8 | 17 | 155 |
| Parahippocampal gyrus | 9 | 21 | 152 |
| Other neocortical regions | 10 | 23 | 277 |
| **Total** | **18** | **43** | **1,233** |

We first sought to determine if these cells fired phase-synchronously with neural oscillations in the ipsilateral hippocampus at rates reliably above chance. For each neuron outside the hippocampus, we used Morlet wavelet convolution to compute spike-coincident hippocampal LFP phases over a broad frequency range (16 log-spaced frequencies from 0.5 to 90.5Hz) across the recording session (25.4 ± 6.9min, M ± SD). We then calculated the mean resultant length (MRL) of the spike-phase distribution at each frequency and Z-scored these values (within-frequency) against null distributions of MRLs from circularly-shifted spike trains. This procedure corrected for spurious contributions to apparent phase-locking, including auto-correlated spiking and nonuniform phase distributions in the hippocampal LFP (Siapas et al., 2005). We refer to the Z-scored MRL as a neuron’s phase-locking strength, which measures the degree to which it fired at a preferential hippocampal LFP phase at a given frequency. Finally, we derived a single phase-locking *p*-value for each neuron by comparing its maximum phase-locking strength across frequencies to the null distribution of maximum phase-locking strengths.

This analysis revealed that 362/1,233 neurons (29.4%) phase-locked to ipsilateral hippocampal LFPs at statistically reliable rates (FDR-corrected across all comparisons with *α* = 0.05). All 18 subjects had at least one significantly phase-locked neuron, and phase-locking rates were 31.5% ± 17.3% (M ± SD) at the subject level. We identified neurons both within and outside the MTL that phase-locked to hippocampal LFPs at varying frequencies and in a highly time-resolved manner (Figure 1B-G). For many neurons, neural oscillations could be seen in the raw hippocampal LFPs (Figure 1A) and in spike-triggered average LFPs that persisted for several cycles surrounding spike onset. This is notable because although hippocampal oscillations in humans occur in short bouts relative to the sustained rodent theta rhythm (Watrous et al., 2013; M. Aghajan et al., 2017), our data suggest that many neurons outside the hippocampus can nonetheless time their firing to coincide with these oscillatory intervals.

**Figure 1:**
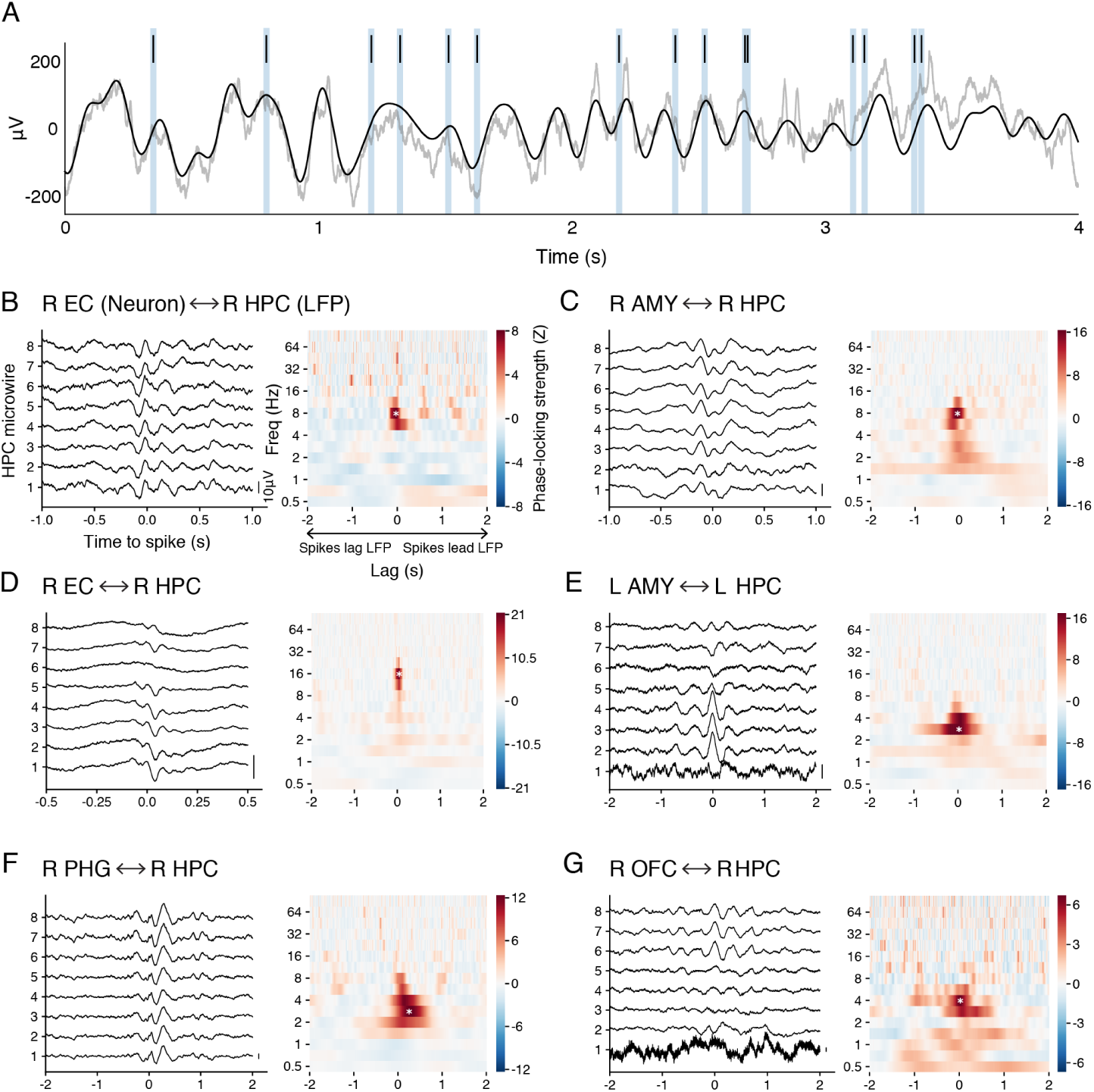
Single-neuron phase-locking examples. (A) A 4s example trace shows spikes from an EC neuron (top) that fired preferentially near the theta peaks of a simultaneously-recorded hippocampal LFP (bottom). The raw hippocampal LFP is shown in light gray, and the same signal, bandpass-filtered from 2-8Hz, is overlaid in dark gray to highlight theta rhythmicity. Panel (B) shows the statistical phase-locking results for this neuron, across the entire recording session. (B-G) Hippocampal phase-locking results are shown for six example neurons from six different subjects. For each neuron, the left graph shows the spike-triggered average LFPs from eight, adjacent micro-electrodes in the hippocampus (these plots are approximately flat in the absence of phase-locking), and the right graph shows the neuron’s permutation test-derived phase-locking strength (Z-score), averaged across these electrodes, as a function of LFP frequency and spike-LFP lag. Asterisks indicate maximum phase-locking strengths. AMY = amygdala; EC = entorhinal cortex; HPC = hippocampus7; L = left; LFP = local field potential; OFC = orbitofrontal cortex; PHG = parahippocampal gyrus; R = right.

One potential explanation for these findings is that phase-locking to hippocampal LFPs occurred as a byproduct of phase-locking to oscillations in a cell’s local vicinity that were themselves phase-synchronous with oscillations in the hippocampus. However, several observations argue that this indirect association cannot solely explain our results. First, if neurons phase-locked to the hippocampus as a consequence of being directly entrained by local oscillations, then they should phase-lock to these local oscillations at least as strongly as they phase-locked to the hippocampus. This is difficult to reconcile with our results, in which 248/362 neurons (68.5%) phase-locked more strongly to hippocampal LFPs than to locally-recorded LFPs at their preferred hippocampal phase-locking frequency (Z = 4.6 ± 3.5, M ± SD difference in phase-locking strength). Moreover, 79/362 neurons (21.8%) phase-locked preferentially to the hippocampus at a frequency at which phase-locking to the local LFP was statistically insignificant. This number on its own greatly exceeds the expected number of false positives (18.1 neurons) under FDR correction. Finally, to directly test whether phase-locking to hippocampal oscillations depended on phase-locking to local oscillations, we employed a spike subsampling technique that removed the influence of local phase-locking on hippocampal phase-locking by enforcing a uniform phase distribution on locally-recorded LFPs (see Methods). This procedure decreased phase-locking strengths to the hippocampus by a significant amount (Z = −1.5 ± 2.1, M ± SD change in phase-locking strength post-correction; *χ*^2^ (1) = 139.6, *p* < 0.001, likelihood ratio test, linear mixed-effects model with nested subject and neuron random effects), yet 334/362 (92.3%) neurons nonetheless remained significantly phase-locked (FDR-corrected at *α* = 0.05). We conclude that our data captured inter-regional interactions between spikes and hippocampal LFPs that occurred largely independently of phase-locking to local oscillations. This suggests that these measures can also differ meaningfully in their relations to memory processes and behavior.

### Phase-locking to the hippocampus by region and frequency

Our observation of widespread synchrony between neural firing outside the hippocampus and rhythmic activity within it offers a means to investigate how the hippocampus is functionally connected to other brain regions at single-neuron resolution. We hypothesized that these functional connections would be especially prevalent in regions that are structurally connected to the hippocampus, and indeed, nearly half of the neurons in the EC (106/221 neurons, 48.0%) and amygdala (161/327, 49.2%) phase-locked to hippocampal LFPs (Figure 2). We also found moderate phase-locking rates in memory-related regions within the extended hippocampal system (Eichenbaum, 2000; Moscovitch et al., 2016), including the parahippocampal gyrus (20/152, 13.2%), orbitofrontal cortex (16/155, 10.3%), and sparsely sampled neocortical regions that included the temporal pole, mid-to-posterior cingulate cortex, and superior temporal cortex (54/277, 19.5%). To assess whether phase-locking rates differed significantly between regions, we fit a logistic mixed-effects model to estimate the probability that a neuron phase-locked to the hippocampus as a function of its region (Table 1) and a random effect of subject. This model confirmed that phase-locking to the hippocampus did not occur uniformly throughout the brain (*χ*^2^(5) = 195.1, *p* < 0.001, likelihood ratio test), as EC and amygdala neurons phase-locked at significantly higher rates than cells in the parahippocampal gyrus, orbitofrontal cortex, anterior cingulate, and remaining neocortical regions (*p* < 0.001, post-hoc pairwise Z-tests, Bonferroni-Holm adjusted). Phase-locking rates did not differ between the EC and amygdala, and we did not test post-hoc comparisons between remaining region pairs.

**Figure 2:**
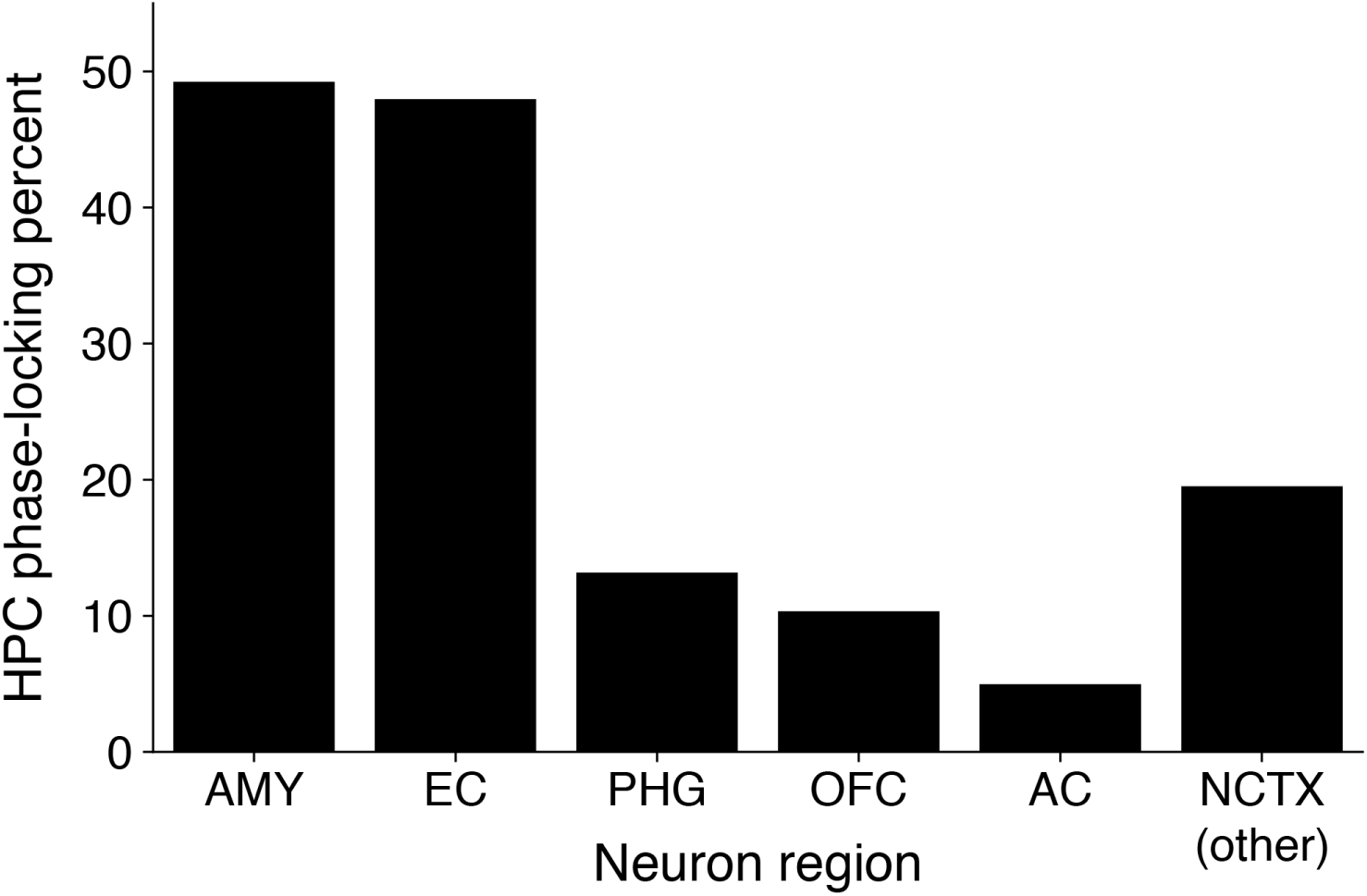
Hippocampal phase-locking rates by region. Each bar shows the percentage of neurons in a given region (pooled across subjects) that phase-locked significantly to ipsilateral hippocampal LFPs at any of the examined frequencies (0.5-90.5Hz). AMY = amygdala; AC = anterior cingulate; EC = entorhinal cortex; HPC = hippocampus; NCTX (other) = sparsely sampled neocortical regions (< 100 neurons or < 5 subjects); OFC = orbitofrontal cortex; PHG = parahippocampal gyrus.

We next sought to determine the frequencies at which phase-locking to the hippocampus occurred. In rodents, neurons outside the hippocampus primarily phase-lock to hippocampal theta oscillations (Kocsis and Vertes, 1992; Siapas et al., 2005; Sirota et al., 2008; Fujisawa and Buzsáki, 2011; Bienvenu et al., 2012; Fernández-Ruiz et al., 2017; Ito et al., 2018), whereas neurons within the hippocampus can phase-lock to both theta and gamma (25-100Hz) rhythms in their local vicinity (Colgin, 2016). In the human hippocampus, neurons have similarly been found to synchronize with both theta and gamma rhythms (Jacobs et al., 2007), while inter-regional phase-locking characteristics are unknown. To address this question, we first plotted the phase-locking strengths at each frequency for all 362 significantly phase-locked neurons. This graph revealed a strong preference for phase-locking to hippocampal LFPs at theta frequencies, with 258/362 (71.3%) neurons being maximally phase-locked from 2-8Hz (Figure 3A). Higher frequency phase-locking was rare, although a small fraction of EC and amygdala neurons phase-locked to beta (16-30Hz) or gamma (>30Hz) frequencies, typically in conjunction with theta entrainment (Figs. 1d and 4a show two examples, from different subjects).

**Figure 3:**
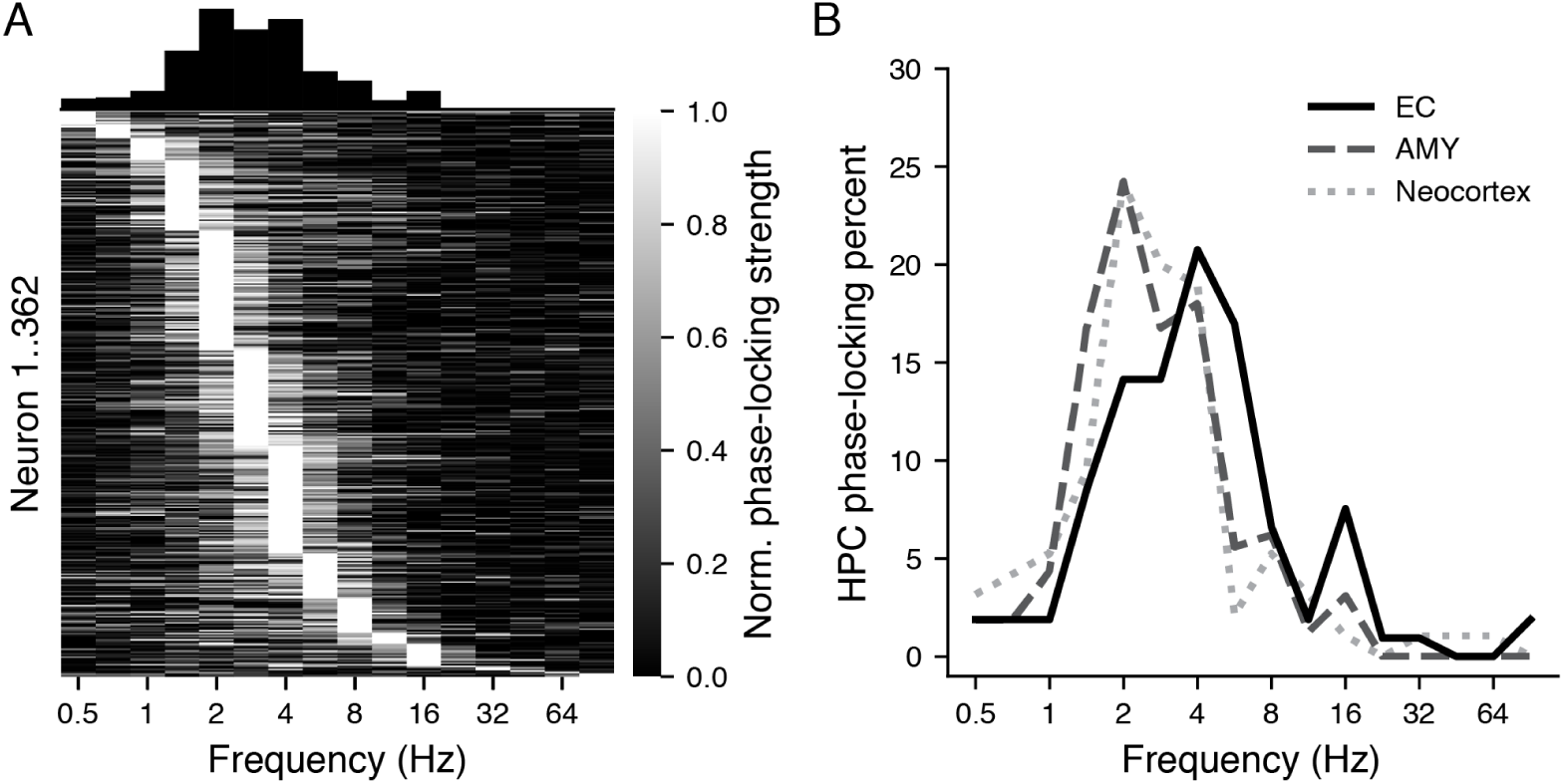
Phase-locking to hippocampal LFPs across frequencies. (A) Normalized phase-locking strengths are shown at 16 frequencies (0.5-90.5Hz) for all 362 neurons that phase-locked significantly to ipsilateral hippocampal LFPs (each row = one neuron). For visual clarity, we divided each neuron’s phase-locking strengths by their maximum value and sorted neurons according to their preferred phase-locking frequency. The distribution of preferred frequencies is shown in the histogram at top. (B) Distributions of preferred hippocampal phase-locking frequencies are shown for all significantly phase-locked neurons in the entorhinal cortex (*n* = 106; solid, black line), amygdala (*n* = 161; dashed, dark gray line), and neocortex (*n* = 95; dotted, light gray line). Y-axis values are normalized such that the area under each curve sums to 100.

Next, we assessed inter-regional differences in the “preferred” frequencies at which neurons phase-locked to the hippocampus (defined by their maximum phase-locking strengths), contrasting cells in the EC (*n* = 106; geometric mean preferred frequency = 4.0Hz), amygdala (*n* = 161; 2.6Hz), and neocortex (*n* = 95; 2.7Hz; aggregating neocortical regions from Table 1 due to the low number of significantly phase-locked cells per region). Preferred phase-locking frequencies differed significantly between these regions (*χ*^2^(2) = 12.5, *p* = 0.002, likelihood ratio test, linear mixed-effects model), with EC neurons phase-locking to the hippocampus at higher frequencies than neurons in the amygdala (Z = 3.0, *p* = 0.007) and neocortex (Z = 3.0, *p* = 0.007), while phase-locking frequencies between the amygdala and neocortex did not differ (Z = 0.3, *p* = 0.758) (post-hoc pairwise Z-tests, Bonferroni-Holm adjusted) (Figure 3B). These inter-regional differences were unchanged in significance when restricting analyses to the 258 theta-phase-locked (2-8Hz) neurons. In summary, neurons outside the hippocampus predominantly phase-locked to hippocampal theta oscillations, and EC neurons phase-locked to a faster theta rhythm than did cells in other regions. To our knowledge, inter-regional differences in hippocampal theta phase-locking frequency have not been previously reported in rodents or other animals.

### Synchronous firing with the contralateral hippocampus

Our finding that many neurons outside the MTL phase-locked to hippocampal LFPs suggests that neurons can synchronize with oscillations across polysynaptic distances. However, tracing studies in monkeys have identified monosynaptic connections between the hippocampus and polymodal association cortices that bypass canonical entorhinal and thalamic relays (Amaral, 1987), so it is possible that the phase-locking effects we observed in neocortical neurons stemmed from such connections. In contrast, neocortical and amygdala neurons have minimal direct connections with the contralateral hippocampus in monkeys (Amaral et al., 1984; Pikkarainen et al., 1999). To gain clearer perspective on the constraints that structural anatomy imposes on inter-regional spike-LFP interactions, we therefore assessed phase-locking to the contralateral hippocampus in 1,057 neurons from 15 subjects with bilateral hippocampal electrodes, employing identical methods as were used to assess ipsilateral phase-locking.

This analysis revealed that 110/1,057 neurons (10.4%) from 13/15 subjects phase-locked significantly to contralateral hippocampal LFPs (FDR-corrected across all comparisons with *α* = 0.05). These cells were broadly distributed throughout the regions that we recorded, although phase-locking to the contralateral hippocampus occurred at considerably lower rates than to the ipsilateral hippocampus (*χ*^2^(1) = 171.9, *p* < 0.001, likelihood ratio test, logistic mixed-effects model with nested subject and neuron random effects). Contralateral phase-locking was essentially restricted to low frequencies, as all but two neurons phase-locked maximally at ≤ 8Hz.

Spike-LFP relations did not emerge independently in each hemisphere; instead, a substantially larger-than-chance fraction of contralaterally phase-locked neurons (84/110, 76.4%) phase-locked significantly to the ipsilateral hippocampus as well (*χ*^2^(1) = 105.4, *p* < 0.001, likelihood ratio test, logistic mixed-effects model). Figure 4A,B shows two representative neurons that exhibited concurrent phase-locking to ipsilateral (top) and contralateral (bottom) hippocampal LFPs. These examples illustrate two differences between ipsilateral and contralateral phase-locking that we observed across the population of 84 bilaterally phase-locked cells. First, ipsilateral phase-locking was stronger than contralateral phase-locking (*χ*^2^(1) = 22.4, *p* < 0.001, likelihood ratio test, linear mixed-effects model with nested subject and neuron random effects) (Figure 4C); and second, ipsilateral phase-locking occurred at higher preferred frequencies (*χ*^2^(1) = 27.4, *p* < 0.001, likelihood ratio test, linear mixed-effects model with nested subject and neuron random effects) (Figure 4D). These variables were significantly correlated, such that neurons with greater decreases in phase-locking strength from ipsilateral to contralateral hippocampus had correspondingly greater decreases in phase-locking frequency (*χ*^2^(1) = 10.4, *p* = 0.001, likelihood ratio test, linear mixed-effects model with nested subject and neuron random effects) (Figure 4E). These results indicate that rhythmic hippocampal activity can propagate between hemispheres and likely at polysynaptic distances, with joint decreases in phase-locking strength and frequency as neurons become farther removed from the hippocampus.

**Figure 4:**
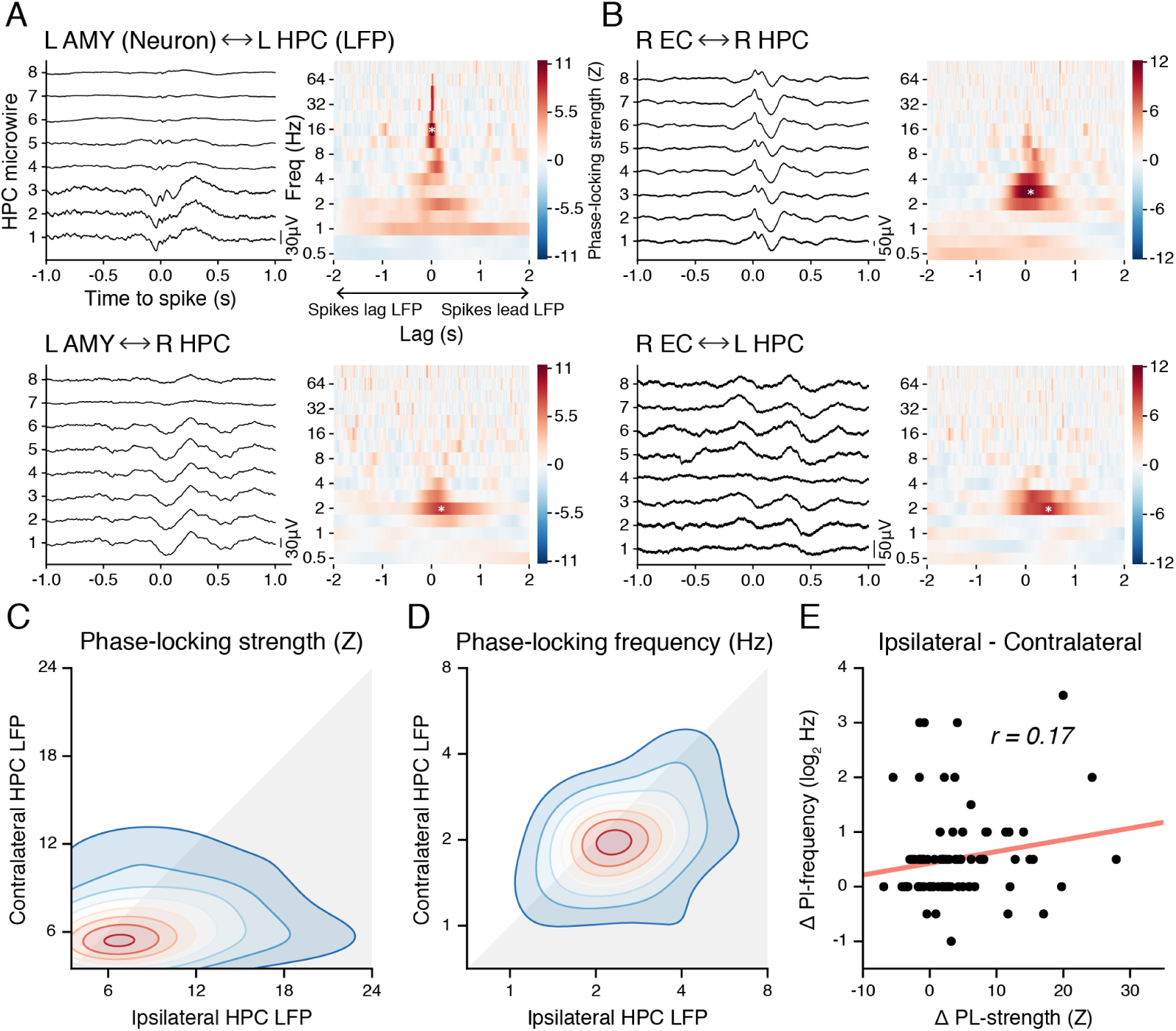
Bilateral phase-locking. (A, B) Spike-triggered LFPs and lag-frequency plots are shown for two example neurons, from different subjects, that phase-locked to both ipsilateral (top) and contralateral (bottom) hippocampal LFPs. (C, D) Gaussian kernel density plots show the paired-sample distributions of maximum phase-locking strengths (C) and preferred phase-locking frequencies (D) to ipsilateral and contralateral hippocampal LFPs, respectively, among 84 bilaterally phase-locked neurons. (E) Differences in the maximum phase-locking strength to ipsilateral versus contralateral hippocampal LFPs are plotted against differences in preferred phase-locking frequencies among 84 bilaterally phase-locked neurons.

### Phase-locking strength fluctuates with changes in hippocampal power

Our findings suggest that phase-locking to hippocampal theta is central to the functional interactions between the hippocampus and multiple other brain regions. One prediction that extends from this interpretation is that when neurons phase-lock strongly to hippocampal theta, there are likely to be co-occurring changes in other aspects of neural activity, beyond theta phase alone. We evaluated this hypothesis by asking whether strong theta phase-locking coincided with distinguishable LFP power patterns in the hippocampus or in a phase-locked neuron’s local region, compared to power during baseline spiking activity (i.e. spikes selected without regard to hippocampal theta phase). Specifically, for the 258 neurons that phase-locked preferentially to ipsilateral hippocampal LFPs at theta frequencies (2-8Hz), we defined the 20% of spikes that were most closely aligned to each neuron’s mean firing phase at its preferred frequency as being “strongly phase-locked” (Figure 5A). We calculated the mean power in the local region and ipsilateral hippocampus, respectively, at 16 frequencies (0.5-90.5Hz) during these strongly phase-locked spikes, and then Z-scored these values against null distributions of mean powers during randomly-drawn spike subsets. Finally, to determine if power changes occurred specifically in these regions or reflected whole-brain dynamics, we performed the same analysis on power values that were averaged over electrodes in all remaining regions.

**Figure 5:**
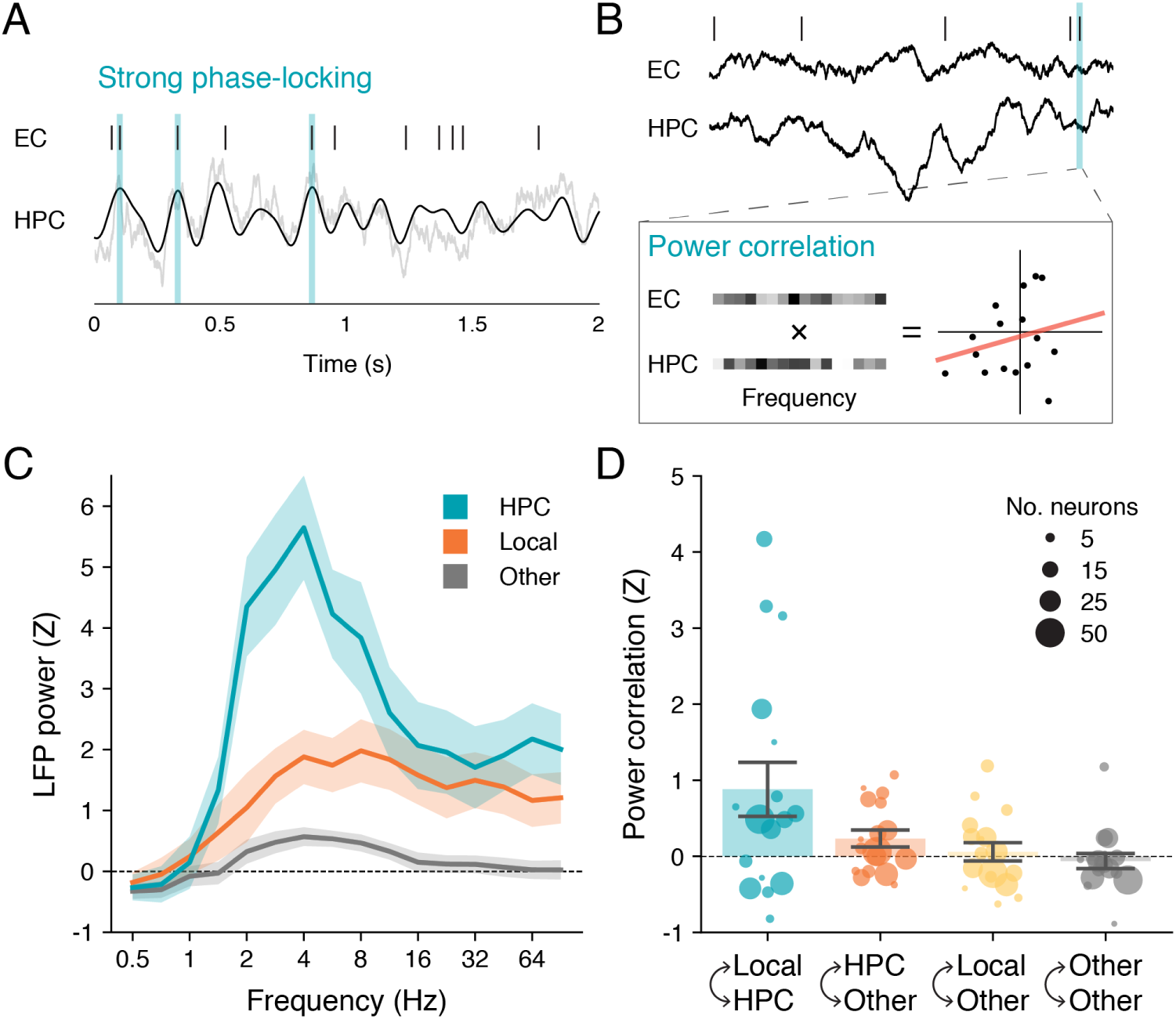
Theta phase-locking relations to hippocampal power and power correlation. (A) Illustrated method for identifying strong phase-locking to hippocampal theta. We selected as “strongly phase-locked” the 20% of spikes across the recording session that were closest to the mean phase at each neuron’s preferred theta frequency. Three such spikes are highlighted in cyan (note the alignment with theta peaks). (B) Illustrated method for calculating power correlations between regions. For each neuron, at each spike time, we calculated the Pearson correlation between LFP powers at 16 frequencies (0.5-90.5Hz; Z-scored within-frequency) in the hippocampus (bottom trace) and in a neuron’s local region (top trace). (C, D) Differences between LFP powers (C) and power correlations (D) are shown during strong phase-locking versus randomly-drawn spike distributions (Z-scores). Values were aggregated across neurons within each subject and then graphed as mean ± SEM across subjects. Each circle in (D) indicates the mean for one subject, with circle area proportional to the number of theta-phase-locked neurons for that subject.

Across neurons, and reliably across subjects, strong phase-locking to hippocampal theta coincided with substantially increased hippocampal theta power (Z = 4.6 ± 2.7, M ± SD for 2-8Hz power across subjects) and a comparatively moderate increase in hippocampal HFA (Z = 1.9 ± 2.4, M ± SD for 32-90.5Hz power across subjects), relative to baseline spiking (Figure 5C, cyan). HFA was also increased in the local region during strong phase-locking (Z = 1.3 ± 1.7, M ± SD across subjects), but in the absence of a similar theta power peak (Figure 5C, orange). In contrast, remaining regions showed minimal to no power changes at any frequency during strong phase-locking (Figure 5C, gray). We confirmed these results by fitting linear mixed-effects models with a single fixed effect (Z-theta power or Z-HFA) and nested subject and neuron random effects. Z-theta power differed significantly between the hippocampus, local region, and remaining regions (*χ*^2^(2) = 362.2, *p* < 0.001, likelihood ratio test), with post-hoc tests revealing greater Z-theta power in the hippocampus than the local region (Z = 15.9, *p* < 0.001) or remaining regions (Z = 22.2, *p* < 0.001), and greater Z-theta power in the local region than remaining regions (Z = 6.3, *p* < 0.001) (pairwise Z-tests, Bonferroni-Holm adjusted). Z-HFA also differed significantly between regions (*χ*^2^(2) = 76.8, *p* < 0.001, likelihood ratio test), with post-hoc tests showing greater Z-HFA in the hippocampus (Z = 8.0, *p* < 0.001) and local region (Z = 7.8, *p* < 0.001) than in remaining regions, and no differences between the hippocampus and local region (Z = 0.2, *p* = 0.848) (pairwise Z-tests, Bonferroni-Holm adjusted).

Lastly, the theory that neural oscillations guide inter-regional communication holds that strong phase-locking to hippocampal theta acts to increase the gain on signaling between the hippocampus and the phase-locked region (Fries, 2005). If this model is correct, we reasoned that strong phase-locking would likely coincide with greater similarity between local and hippocampal neural activity patterns as a consequence of their increased interactions. To test this hypothesis, we examined neural similarity patterns in the spectral power domain, consistent with prior intracranial EEG studies in humans (Manning et al., 2011; Staresina et al., 2016). Specifically, for each of the 258 theta-phase-locked neurons, we calculated the Pearson correlation at each spike time between LFP powers across 16 frequencies (0.5-90.5Hz; Z-scored within frequency) in the local region and powers at the corresponding frequencies in the ipsilateral hippocampus (Figure 5B). For each neuron, we then computed the mean power correlation during strongly phase-locked spikes and Z-scored this value against a null distribution of mean power correlations during randomly-drawn spike subsets. To determine if power correlation changes occurred specifically between the local region and hippocampus, we repeated this analysis for three additional categories of inter-regional pairs: (1) connections between the ipsilateral hippocampus and all regions except the local region; (2) connections between the local region and all regions except the ipsilateral hippocampus; and (3) connections between all remaining regions.

Consistent with our hypothesis, power correlations between the local region and hippocampus were reliably increased during strong phase-locking compared to baseline spiking (Z = 0.9 ± 1.4, M ± SD across subjects) (Figure 5D, cyan). In contrast, power correlations differed minimally in the remaining three categories of inter-regional pairs (Figure 5D, orange, yellow, and gray). We confirmed these results by fitting a linear mixed-effects model with a fixed effect of Z-scored power correlation and nested subject and neuron random effects. Z-power correlations differed significantly between connection categories (*χ*^2^(3) = 80.9, *p* < 0.001, likelihood ratio test), with local-to-hippocampal connections having higher Z-power correlations than each of the three remaining categories (all *p* < 0.001, post-hoc pairwise Z-tests, Bonferroni-Holm adjusted), and no significant differences between the remaining category pairs.

## Discussion

We asked whether single neurons outside the hippocampus fired phase-synchronously with hippocampal neural rhythms in the human brain while subjects (18 neurosurgical patients) played a virtual navigation game. Confirming evidence from many studies in rodents (Siapas et al., 2005; Kocsis and Vertes, 1992; Sirota et al., 2008; Fujisawa and Buzsáki, 2011; Bienvenu et al., 2012; Fernández-Ruiz et al., 2017; Ito et al., 2018), we found that a substantial fraction of extrahippocampal neurons (nearly 30% of 1,233 recorded cells) exhibited significant phase-locking to hippocampal LFPs, principally within the theta frequency range.

Prior studies have assessed functional connections between memory-related regions using implanted macroelectrodes, which integrate activity over many thousands of neurons (Babiloni et al., 2009; Watrous et al., 2013; Solomon et al., 2017; Zheng et al., 2019; Kunz et al., 2019). Our results extend these findings to demonstrate that cortico-hippocampal functional connections also emerge at the single-neuron level (Figure 1), and despite orders of magnitude separating micro- and macro-recording scales, both approaches highlight a privileged role for theta synchronization during memory-dependent tasks (Figure 3). Phase-locking to the hippocampus was most common among neurons in structurally-connected regions (the EC and amygdala) but also occurred in more remote, neocortical regions (Figure 2), as well as to the contralateral hippocampus (Figure 4). These findings parallel results from noninvasive neuroimaging analyses in healthy adults, in which functional MRI connectivity patterns have been shown to overlap and exceed the limits of white matter tractography (Honey et al., 2009). Finally, we observed regionally-specific LFP power changes when neurons fired near their preferred hippocampal theta phase (“strong phase-locking”), relative to baseline spiking. Strong theta phase-locking coincided with increased hippocampal theta power (Figure 5C, cyan), greater local and hippocampal HFA (Figure 5C, cyan and orange), and increased cross-frequency power correlations between the hippocampus and a phase-locked neuron’s local region (Figure 5D, cyan). These results confirm the hypothesis that hippocampal theta rhythms coordinate the timing of single-neuron activity throughout the human episodic memory system. Such coordination could facilitate precise interactions between the hippocampus and other brain regions to determine the content and quality of episodic memories.

Our results support several predictions of the “Communication through Coherence” (CTC) hypothesis, a leading model of oscillatory involvement in inter-regional communication (Fries, 2005). According to this hypothesis, for oscillations to enhance the effective coupling between two regions, neurons in both regions must phase-lock to the same rhythm (Fries, 2005). While previous studies identified neurons within the human hippocampus that phase-locked to local theta oscillations (Jacobs et al., 2007; Rutishauser et al., 2010; Watrous et al., 2018; Kamiński et al., 2020), here we established that neurons in many additional regions also phase-lock to this rhythm (Figure 2). Notably, although hippocampal theta in humans usually occurs in short, ∼1-2s “bouts” of elevated power (Watrous et al., 2013; M. Aghajan et al., 2017), both the extrahippocampal neurons in our study and hippocampal neurons in the study by Jacobs et al. (2007) exhibited strongest phase-locking during high theta power intervals (Figure 5C). These results suggest that high hippocampal theta power reflects increased engagement between the hippocampus and other memory-related regions, despite that hippocampal theta power has shown inconsistent correlations with memory performance (Herweg et al., 2020).

Next, the CTC hypothesis states that oscillations can increase the effective coupling between regions by synchronizing spike times in one region with the phases at which postsynaptic neurons in another are most excitable, producing selective input gain (Fries, 2005). This process induces transiently greater activity in both regions if these postsynaptic neurons are successfully discharged (Fell and Axmacher, 2011). Our finding that strong phase-locking to hippocampal theta coincided with greater HFA both locally and in the hippocampus suggests that phase-locking can have such an effect (Figure 5C), as HFA is closely correlated with aggregated, local neural firing levels (Manning et al., 2009; Nir et al., 2007).

Finally, the consequence of optimal phase-locking, according to CTC, is a transient increase in the information shared between two regions (Fries, 2005). Our results offer preliminary support for this hypothesis in showing that strong phase-locking to hippocampal theta coincided with increased power correlations between the local region and hippocampus (Figure 5D). Importantly, this effect occurred selectively between these regions, as power correlations were not globally increased between the hippocampus and other regions all at the same time. Stronger support for this hypothesis would come from comparing hippocampal theta phase-locking to regional activity correlations with behavioral stimuli, an analysis that might also help to delineate the direction of information transfer between the hippocampus and other regions under varying task conditions.

A fundamental question concerns the frequency range over which human hippocampal theta occurs – which, due to the absence of continuous theta oscillations in humans and the diversity of analytical approaches that have been used to study them, has produced disagreement in the field (Herweg et al., 2020; Jacobs, 2014). Most studies have approached this question from a behavioral angle and have reported a wide mixture of findings; for example, some studies identified hippocampal theta effects in the ∼4-10Hz (“rodent theta”) range that correlated with navigation and memory performance (Fell et al., 2011; M. Aghajan et al., 2017; Bohbot et al., 2017), others found effects on similar tasks but at slower, ∼1-4Hz frequencies (Watrous et al., 2013; Jacobs, 2014; Lega et al., 2016; Kunz et al., 2019), and still others found effects within both of these frequency ranges (Lega et al., 2012; Bush et al., 2017; Zheng et al., 2019), suggesting that hippocampal theta in humans either spans a very broad range or else refers to two, poorly-differentiated rhythms that might serve different functions.

In this study, by focusing on neurophysiological and network-based analyses, we offer a complementary perspective on how hippocampal theta in humans can be viewed. We interpret our results together with three principle claims. First, hippocampal theta appears to be slower in humans than in rodents (also see Jacobs, 2014). Although a small number of neurons in our dataset phase-locked to frequencies typical of rodent theta (Figure 1B,C), a much larger fraction of cells fired synchronously with 2-4Hz hippocampal rhythms (Figure 3A), and this effect was consistent across subjects and brain regions (Figure 3B).

Second, slower theta frequencies appear to support weak interactions with regions farther removed from the hippocampus. The presence of neo-cortical and contralateral phase-locking suggests that theta oscillations can propagate polysynaptically, consistent with their acting as traveling waves across the hippocampus and neocortex (Zhang et al., 2018). However, the temporal precision between spikes and hippocampal theta phase would likely decrease at greater synaptic path lengths, favoring weaker phase-locking to slower frequencies that can better accommodate spike-time variability. This reasoning is supported by our finding: (1) that compared to EC neurons, neocortical cells were both substantially less likely to phase-lock to the hippocampus (Figure 2) and preferred slower theta frequencies (Figure 3B); (2) that among bilaterally phase-locked neurons, phase-locking to the contralateral hippocampus was weaker (Figure 4C) and slower (Figure 4D) than to the ipsilateral hippocampus; and (3) changes in phase-locking strength and frequency from ipsilateral to contralateral hippocampus were significantly correlated across neurons (Figure 4E). Theta frequency variability could thus reflect a trade-off between how precisely the hippocampus can coordinate its activity with other regions (preferring faster theta) and the spatial extent over which this coordination occurs (preferring slower theta). This might further explain why the neurons we recorded outside the hippocampus rarely phase-locked to the hippocampal beta or gamma oscillations (Figure 3A), despite an earlier study finding that neurons within the human hippocampus commonly phase-locked to these fast oscillations (Jacobs et al., 2007).

Third, theta frequency relations likely vary by location along the hippocampal longitudinal axis, with faster theta supporting functional interactions with the posterior hippocampal system. This claim is supported by our finding that amygdala neurons, which project primarily to the anterior hippocampus, phase-locked to slower hippocampal theta frequencies than did EC neurons, which project more evenly throughout the longitudinal extent of the hippocampus (Figure 3B) (Strange et al., 2014). Faster theta oscillations were also found in the posterior than anterior hippocampus by a study that directly compared electrodes between these regions (Goyal et al., 2020). Taken together, our results suggest that hippocampal theta in humans is not a single oscillation but instead encompasses a range of low frequency activity, centered around ∼4Hz, that is best understood in context with the functional interactions between the hippocampus and its extended network. We hypothesize that the “dominant” theta frequency (i.e. the frequency with the most power) changes over time as these interactions are reorganized to aid communication with different combinations of regions. Viewed from this perspective, the frequency instability of hippocampal theta in humans compared to rodents could result from the expanded neocortex and increased complexity of cortico-hippocampal interactions in higher-order mammals.

This study has several important limitations. First, all subjects had pharmacoresistant epilepsy, and we cannot rule out that some results might stem from pathological activity. We are encouraged by the general agreement between our results and those reported in rodents (Siapas et al., 2005; Kocsis and Vertes, 1992; Sirota et al., 2008; Fujisawa and Buzsáki, 2011; Bienvenu et al., 2012; Fernández-Ruiz et al., 2017; Ito et al., 2018), but clearer evidence that phase-locking to hippocampal theta reflects normal physiology in humans will come from comparing this measure to memory performance. Our study features a large sample size relative to most singleneuron studies in humans, but the virtual navigation task that was used had minimal memory requirements (Ekstrom et al., 2003), and we decided against analyzing other behavioral contrasts for which null results would have been difficult to interpret. A second limitation concerns the quality of single-neuron isolation in this study. We recorded spikes from singlemicrowires with limited ability to resolve distinct spike waveforms, and so it is likely that some “single-neurons” in our study combine spikes from a small number of neighboring cells. However, to the extent that this issue affected our analyses, it would presumably interfere with the ability to detect significant phase-locking to the hippocampus, yielding an underestimate of phase-locking rates rather than an excess of false positive results.

Still little is known about the relations between theta phase-locking and human episodic memory, but early results suggest an exciting future for this field. Prior studies have focused on the behavioral correlates of phase-locking to local theta rhythms within the MTL; for example, successful image encoding was found to depend on theta phase-locking strength among hippocampal and amygdala neurons (Rutishauser et al., 2010), while an-other study found that MTL neurons can represent contextual information in their theta firing phase (Watrous et al., 2018). Here we showed that hippocampal theta oscillations also inform the timing of widespread cortical activity, and these inter-regional relations existed largely independently of phase-locking to local theta rhythms. From a physiological vantage, theta oscillations thus appear to be positioned at the interplay between local and inter-regional neural computations in the human memory system. Going forward, it will be especially interesting to consider how local and inter-regional phase-locking effects can be teased apart behaviorally, and whether these measures offer independent or overlapping predictions of memory performance. Such analyses could be well positioned to unite findings from animal and human studies and advance a more mechanistic account of episodic memory organization across multiple levels of scale, from single neurons to macroscopic fields.

## Methods

### Participants

Subjects were 18 patients with pharmacoresistant epilepsy who were implanted with depth electrodes to monitor seizure activity. Clinical teams determined the location and number of implanted electrodes in each patient. We performed bedside cognitive testing on a laptop computer in consenting patients. The experiment was approved by institutional review boards at the University of California, Los Angeles and the University of Pennsylvania.

### Spatial navigation task

We analyzed data from 43 testing sessions (1-4 sessions per subject, median = 2.5) that were each 25.4 ± 6.9min (M ± SD) in length. During each session, subjects played a first-person, spatial navigation game called *Yellow Cab*, in which they drove freely around a virtual town while alternating between searching for passengers who appeared at random locations and dropping them off at their requested locations. The specifics of this task along with subjects’ performance have been described previously in detail (Ekstrom et al., 2003).

### Recording equipment

Each subject was implanted with six to 12 Behnke-Fried depth electrodes that feature macroelectrode contacts for clinical monitoring and 40µm diameter, platinum-iridium microwires for measuring microscale local field potentials (LFPs) and extracellular action potentials (Fried et al., 1999). Electrode localizations were confirmed by the clinical team using post-operative structural MRIs or post-operative CT scans that were co-registered to pre-operative structural MRIs. Microwires were packaged in bundles of eight high-impedance recording wires and one low-impedance wire that served as the recording reference. Each microwire bundle was threaded through the center of a depth probe and extended 5mm from the implanted end. Microwire LFPs were amplified and sampled at 28-32kHz on a Neuralynx Cheetah (Neuralynx, Tucson, AZ) or Blackrock NeuroPort (Blackrock Microsystems, Salt Lake City, UT) recording system.

### Spike-sorting

We performed semi-automatic spike sorting and quality inspection on each extrahippocampal microwire channel using the WaveClus software package in Matlab (Quiroga et al., 2004), as previously described (Ekstrom et al., 2003). We isolated 0-4 single-units from each microwire electrode (0.6 ± 0.8, M ± SD). Repeated testing sessions occurred on different days, and we spike-sorted and analyzed these data separately.

### Spectral feature extraction

Hippocampal microwire LFPs were manually checked for recording quality, and we removed 2/35 hippocampal microwire bundles from the analysis due to excessive signal contamination. For the remaining electrodes, we downsampled microwire LFPs to 2kHz, notch-filtered the signal at 60Hz and harmonic frequencies to remove electrical line noise, and then convolved LFP traces with five-cycle Morlet wavelets to extract spectral power and phase at 16, log-spaced frequencies from 0.5 to 90.5Hz (*f* = 2^(*x/*2)−1^Hz for *x* ∈ 0, 1, …, 15). Power values were log-transformed and then Z-scored, within-frequency, across all timepoints in the session, separately for each microwire channel.

### Hippocampal phase-locking strength and significance

To determine if a neuron phase-locked to LFPs from a given hippocampal microwire bundle, we computed the mean resultant length (MRL) of spectral phase values at each spike time during the recording session, for each microwire channel and frequency. The MRL is calculated by dividing the sum across phase-angle unit vectors by the total number of spikes, yielding a measure from zero to one that indicates the extent to which the spike-phase distribution is clustered around a unimodal peak. An MRL = 1 means that all spike times occurred at identical hippocampal LFP phases, while chance association between spikes and LFPs is expected to produce a distribution that approaches uniformity (MRL = 0) at increasing numbers of spikes.

Several factors other than direct spike-LFP interactions can artificially inflate the MRL – for example, if the underlying LFP phase distribution is nonuniform, or if a neuron’s spike train is highly auto-correlated (Siapas et al., 2005)). We used a permutation-based procedure to estimate a neuron’s true phase-locking strength while controlling for these spurious effects. This procedure entailed circularly shifting each neuron’s spike train 1,000 times by random, independent intervals up to the length of the recording session, generating a null distribution of 1,000 MRLs for each microwire channel and frequency. Phase-locking strengths were calculated by averaging MRLs across channels and then Z-scoring a neuron’s actual MRL at each frequency by corresponding values from the null distribution.

To determine which neurons phase-locked significantly to the hippocampus, we obtained a phase-locking *p*-value for each neuron by comparing its maximum phase-locking strength (across frequencies) to the null distribution of 1,000 maximum phase-locking strengths from circularly-shifted spike trains (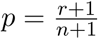, where *r* is the number of replicate values ≥ the actual value, and *n* is the number of replicates (North et al., 2002)). Lastly, we false discovery rate (FDR)-corrected these phase-locking *p*-values at the study level (*n* = 1, 233 neurons), setting *α* = 0.05, using the adaptive linear step-up procedure, which controls the expected proportion of true null hypotheses among rejected nulls for both independent and positively-dependent test statistics, and has greater statistical power than the commonly used Benjamini-Hochberg procedure ((Benjamini et al., 2006)). This correction was applied separately for ipsilateral and contralateral phase-locking comparisons.

### Spike subsampling procedure to remove local phase-locking effects

We sought to determine if inter-regional phase-locking to hippocampal LFPs could be parsimoniously explained by the presence of spike-LFP phase-locking to locally-recorded signals that exhibited LFP-LFP phase synchrony with the hippocampus. To resolve this question, we subsampled spikes from each neuron in a manner that explicitly removed local phase-locking effects while retaining the largest possible number of spikes for reanalysis. We then reassessed phase-locking significance to the hippocampus in the subsampled spike distributions.

The details of this method were as follows: (1) For each neuron that phase-locked significantly to the hippocampus (see “Hippocampal phase-locking strength and significance”), we identified the frequency with the maximum phase-locking strength as the neuron’s “preferred” phase-locking frequency. (2) We calculated spike-coincident LFP phases at the preferred frequency for each of seven neighboring electrodes in the neuron’s microwire bundle. Data from the neuron’s own recording wire were excluded due to spike artifacts in the LFP that can contaminate phase-locking analyses (Jacobs et al., 2007). (3) For each of the resulting spike-phase distributions, we categorized spikes according to which of six, evenly-spaced phase bins (0 to 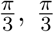 to 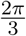, etc.) they corresponded to. We identified the phase bin with the smallest number of categorized spikes, *n*_*min*_, and randomly selected *n*_*min*_ spikes from each of the six phase bins. This process enforced an approximately uniform phase distribution onto the local LFP, and it yielded seven distinct spike subsamples (one for each local electrode). (4) We randomly assigned these spike subsamples to seven (out of eight) hippocampal LFPs. Then, following the method described in our original phase-locking analysis, we quantified the phase-locking strength at the preferred frequency by calculating the mean MRL across spike sub-samples and Z-scoring this value against a null distribution of MRLs from 1,000 circularly-shifted spike trains. A phase-locking *p*-value was calculated as described previously. (5) Finally, *p*-values were FDR-corrected at the study level using the stringent Benjamini-Hochberg procedure (Benjamini et al., 2006).

### LFP power during strongly phase-locked spikes

We examined whether strong phase-locking to hippocampal theta was associated with significant changes in LFP power, a widely studied measure of neural activity. This analysis was performed in the 258 neurons that phase-locked significantly to ipsilateral hippocampal LFPs at a preferred frequency between 2-8Hz, according to the following procedure. First, we defined “strong phase-locking” as the 20% of spikes that were closest in alignment (absolute value of the phase difference) to each neuron’s mean firing phase at its preferred hippocampal theta phase-locking frequency. We then calculated the mean LFP power at each of 16 frequencies from 0.5 to 90.5Hz (see “Spectral feature extraction”) across these strongly phase-locked spikes times, separately for each microwire recording electrode in the brain. For each neuron, this step yielded a frequency × electrode matrix of LFP powers during strong phase-locking. We then calculated the mean power at each frequency across electrodes: (1) in the ipsilateral hippocampus; (2) in a neuron’s local region (electrodes from the same microwire bundle, excluding the neuron’s own recording wire); and (3) in all remaining regions. For each neuron, this step yielded a vector of mean LFP powers during strongly phase-locked firing in each of the three regions-of-interest (hippocampus, local region, and remaining regions).

We next sought to determine if the LFP powers during strong phase-locking differed from those during average spiking activity (i.e. spikes selected without regard to hippocampal theta phase). To estimate LFP power during average spiking, we repeatedly selected 20% of spikes from each neuron at random (1,000 permutations) and calculated the mean power in the hippocampus, local region, and remaining regions following the procedure described above. We then Z-scored the LFP powers during strong phase-locking against those from the randomly drawn spike distributions, separately for each frequency and region-of-interest. Finally, to assess whether there were any reliable differences in LFP power during strong phase-locking compared to average spiking across subjects, we calculated the mean Z-scored power for each frequency and region-of-interest, first across all neurons within each subject, and then across all subjects. The means and standard errors of the resulting values are shown in Figure 5C.

### Cross-frequency power correlations during strongly phase-locked spikes

For each of the 258 theta-phase-locked neurons (see “LFP power during strongly phase-locked spikes”), we calculated cross-frequency power correlations between all pairs of regions, at each spike time, according to the following method. First, LFP powers at each of 16 frequencies (0.5-90.5Hz) were averaged across microwire electrodes in each region (i.e. electrodes within each microwire bundle; for the local region, we excluded the neuron’s own recording electrode), yielding a region × frequency × timepoint array of power values. Next, for each pair of regions, we calculated the cross-frequency power correlation at each spike time as the Pearson correlation between LFP powers across frequencies in one region and powers at the corresponding frequencies in the second region. It is important to note that these power values were log-transformed and Z-scored across session time prior to performing this analysis (see “Spectral feature extraction”), so there was no 1/f decay that would cause lower frequencies to exert an outsized influence on the cross-frequency power correlation. This measure can instead be interpreted as reflecting the instantaneous similarity between activity in two regions in the spectral power domain. This method is similar to representational similarity analyses that have been used in prior intracranial EEG studies that compared LFP powers at two timepoints within the same region, e.g. during the encoding and recall of word pairs (Manning et al., 2011; Yaffe et al., 2014; Staresina et al., 2016). Here we asked whether the similarity between two regions at the same timepoint was influenced by hippocampal theta phase-locking, independent of a behavioral contrast.

To test the hypothesis that strong phase-locking to hippocampal theta co-occurs with increased cross-frequency power correlations, we first calculated the mean cross-frequency power correlation between each pair of regions across strongly phase-locked spikes (see “LFP power during strongly phase-locked spikes”), yielding a vector of power correlations by region-pair. We then further averaged the power correlations across region-pairs within each of four categories: (1) the local region to ipsilateral hippocampus (a single region-pair); (2) the ipsilateral hippocampus to all remaining regions; (3) the local region to all remaining regions; and (4) all region pairs that did not include the local region or ipsilateral hippocampus.

To determine if cross-frequency power correlations differed during strong phase-locking relative to average spiking activity, we employed the same approach as we used to evaluate differences in LFP powers. Specifically, we Z-scored cross-frequency power correlations during strong phase-locking against randomly drawn spike distributions (1,000 permutations, each with 20% of spikes). This procedure resulted in a Z-scored value for each neuron that reflected the degree to which the cross-frequency power correlation differed during strong phase-locking relative to randomly-drawn spike subsets. Finally, to assess whether these Z-scored power correlations differed reliably across subjects, we calculated the mean Z-scored power correlation within each category of region pairs (local-hippocampus, hippocampus-other, local-other, other-other), first across all neurons within each subject, and then across all subjects. Figure 5D shows the means and standard errors of the resulting values (bar plots), along with the means for each subject scaled by the number of neurons (filled circles).

### Statistics

Linear and logistic mixed-effects models with fixed slopes and random intercepts were performed using the lme4 package in R (Baayen et al., 2008). Likelihood ratio tests between full and reduced model fits were used to obtain *p*-values for interpreting significance. All models included a single random effect of subject or, for data in which multiple observations were sampled from each neuron (noted in the Results), nested random effects of subject and neuron identity. In addition, each model included a single fixed effect that was dropped to generate the reduced model. We adopted this approach to control for subject-level differences in our data that would be overlooked when using conventional methods, like linear regression, that interpret each neuron as being independent of all other neurons. This analytical approach was particularly important for comparing effects between regions, because each subject had electrodes placed in only a subset of the regions that we analyzed. For models in which the independent variable was a categorical measure with three or more levels, if the likelihood ratio test revealed a significant effect (*p* < 0.05), we performed post-hoc, pairwise Z-tests on the fitted model terms with Bonferonni-Holm correction for multiple comparisons where noted in the Results.

### Software

Mixed-effects effects models were fit using the lme4 package in R (Baayen et al., 2008). Spike-sorting was performed using the WaveClus software package in Matlab (Quiroga et al., 2004). All additional analyses were performed using code that was developed in-house in Python 3, utilizing standard libraries and the following, publicly-available packages: astropy, mne, numpy, pandas, pycircstat, seaborn, scipy, and statsmodels.

### Data availability

The data used in this study is publicly available for download from the Cognitive Electrophysiology Data Portal (http://memory.psych.upenn.edu/Electrophysiological_Data). This dataset includes de-identified, raw EEG data; spike-sorted single-unit data; and pre-processed phase-locking data. The download also contains code to reproduce our analyses, including a Jupyter notebook that will generate the main table and figure plots, and an R script to run statistical tests.

## Acknowledgments

We are grateful to the patients for their participation and thank hospital staff and researchers who were involved in data acquisition. This work was supported by the National Science Foundation GRFP grant (D.R.S.), NIH U01 (NS113198 to M.J.K.) and NINDS (NS033221 and NS084017 to I.F.), and Deutsche Forschungsgemeinschaft (DFG) Grant HE 8302/1-1 (N.A.H.).

## Author Contributions

D.R.S. and M.J.K. designed the experiment. IF performed surgical procedures and supervised recordings and data collection. D.R.S. analyzed the data. D.R.S. wrote the manuscript with feedback from all authors.

